# More efficacious drugs lead to harder selective sweeps in the evolution of drug resistance in HIV-1

**DOI:** 10.1101/024109

**Authors:** Alison F. Feder, Soo-Yon Rhee, Robert W. Shafer, Dmitri A. Petrov, Pleuni S. Pennings

## Abstract

In the early days of HIV treatment, drug resistance occurred rapidly and predictably in all patients, but under modern treatments, resistance arises slowly, if at all. The probability of resistance should be controlled by the rate of generation of resistant mutations. If many adaptive mutations arise simultaneously, then adaptation proceeds by soft selective sweeps in which multiple adaptive mutations spread concomitantly, but if adaptive mutations occur rarely in the population, then a single adaptive mutation should spread alone in a hard selective sweep. Here we use 6,717 HIV-1 consensus sequences from patients treated with first-line therapies between 1989 and 2013 to confirm that the transition from fast to slow evolution of drug resistance was indeed accompanied with the expected transition from soft to hard selective sweeps. This suggests more generally that evolution proceeds via hard sweeps if resistance is unlikely and via soft sweeps if it is likely.

## 1 Introduction

In the first two decades of the HIV epidemic, HIV became a prime example of fast evolutionary change, especially because of the evolution of drug resistance quickly after initiation of treatment. Nowadays, HIV treatments are more clinically efficacious and the evolution of drug resistance has become much slower and often does not occur for years if at all. The rate at which evolution occurs has been the subject of considerable recent interest in the evotionary biology community. Though traditionally evolution was thought to be slow [1], there are a growing number of examples of fast evolution to selective pressures such as pesticides [2–5], industrialization [6], or antibiotics [7, 8]. HIV represents an interesting case because its evolutionary speed has changed drastically over time.

Population genetic theory suggests that the primary difference between slow and fast evolving populations is driven by the availability of adaptive mutations. In a large population with a high mutation rate, mutations may be available as standing genetic variation (pre-existing variation) or be generated anew every generation, allowing the population to adapt to its environment rapidly. If adaptive mutations are rare, because the population is small, the mutation rate is low, or only few specific mutations (or combinations of mutations) can help a population adapt, the population will likely adapt to its environment much more slowly.

The availability of adaptive mutations does not only change the rate of adaptation, it also changes how adaptation affects genetic diversity in a population. If adaptive mutations are rare, i.e., less then one adaptive mutation occurs per generation in the population, the first successful mutation is likely to rise to high frequency before any subsequent adaptive mutations reach appreciable frequencies (see Figure 1A). This results in a hard selective sweep, in which the single adaptive mutation and the nearby linked mutations becomes fixed in the population (Figure 1B). Hard selective sweeps sharply reduce genetic diversity in the population (Figure 1C) [9, 10] in a similar manner to a strong genetic bottleneck.

**Figure 1:**
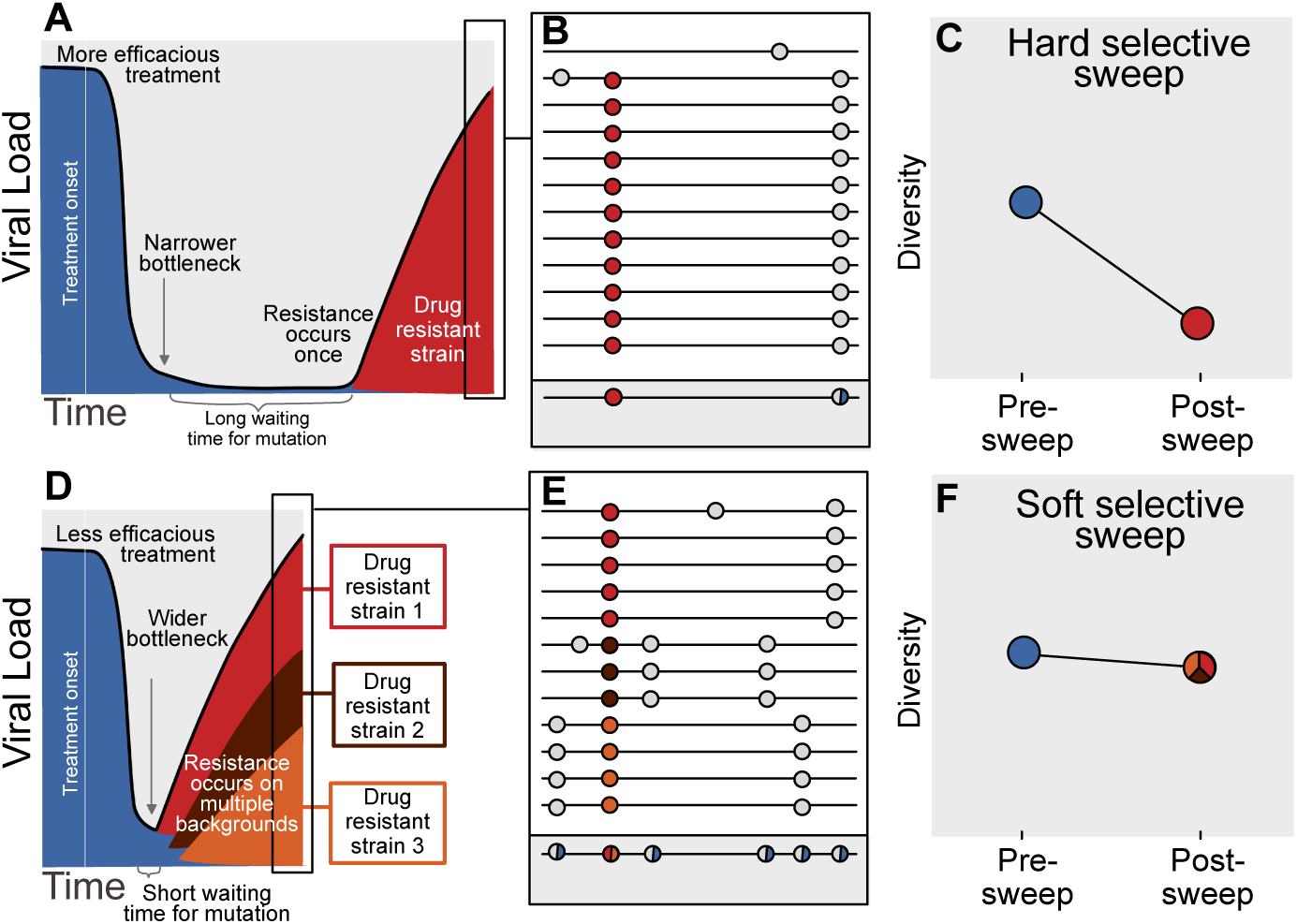
Prediction of drug resistance acquisition with more and less efficacious treatments. Among patients treated with more efficacious treatments (top), we predict HIV populations to have a lower probability of acquiring resistance per generation. As a result, the population must wait a long time for a beneficial genotype, so when resistance does occur, it will spread through the population in a hard selective sweep before other resistant genotypes emerge **(A)**. Since resistance only occurs on a single genetic background (background mutations in grey), all sequences with resistance will be similar **(B)** and diversity following this type of selective sweep will be reduced **(C)**. We can use the reduction of diversity to determine that a selective sweep is hard. In patients treated with less efficacious treatments (bottom), we predict HIV populations should have a higher probability of acquiring resistance per generation, so resistance will be acquired more quickly and selective sweeps of drug resistance mutations will be soft **(D)**. We can detect these soft selective sweeps, because diversity remains high when resistance mutations on different genetic backgrounds rise in frequency simultaneously **(EF)**.

In contrast, when adaptive mutations are common, i.e., more than one occurs per generation in the population, the same adaptive mutation may occur several times in a very short time span on different genetic backgrounds. These adaptive mutations can increase in frequency virtually simultaneously (Figure 1D) [11] and multiple genetic backgrounds are therefore expected to reach substantial frequencies with no single genetic background dominating the population (Figure 1E). This pattern is known as a soft selective sweep and is expected to lead to almost no reduction of genetic diversity (Figure 1F) [12], comparable to a mild bottleneck.

In HIV, the evolution of drug resistance was fast in patients on early anti-retroviral therapies [13], but current multi-drug regimens have substantially slowed the rate of evolution of resistance [14]. In this study, we ask whether this transition from fast evolution of drug resistance in the early years of antiretroviral treatments, to a slow rate of evolution of resistance in patients on more clinically efficacious treatments is associated with a transition from soft selective sweeps to hard selective sweeps. To test whether this is the case, we look at the relationship between fixed drug resistance mutations (DRMs) and genetic diversity across 29 common anti-retroviral drug regimens. The expectation is that when hard selective sweeps predominate, we will find a negative correlation between the number of DRMs and genetic diversity in a population. On the other hand, when soft selective sweeps predominate, we expect to find no such correlation. We use 6,717 HIV sequences from the same number of patients from the Stanford HIV Database [15]. These sequences contain information about the number of DRMs and, as we will further explain in the next paragraph, they also contain information about genetic diversity in the viral population.

Most sequencing of HIV populations in patients is done with the intent to discover DRMs for diagnostic and therapeutic reasons [16]. As such, in standard clinical practice, a sample from a patient’s entire HIV population is amplified via PCR and then sequenced using the traditional Sanger method resulting in a single consensus sequence. Genetic diversity may result in ambiguous calls (also referred to as mixtures) in the reported sequence, so that a signal of within-patient genetic diversity is retained even though there is only a single sequence per patient. We use the ambiguous calls to quantify within-patient genetic diversity (see Figure 1BE, grey box), following several other studies [17–20]. Although ambiguous calls are an imperfect measure of diversity, it has been shown that the signal from ambiguous calls can be reproduced between laboratories [21]. By using ambiguous calls as a proxy for diversity, we are able to take advantage of a large number of HIV-1 sequences, allowing us to study the evolutionary dynamics of HIV drug resistance evolution in a historical perspective[15].

Through examining HIV sequences of 6,717 patients over the past two decades in the presence of many different drug regimens, all sequenced using Sanger sequencing technology, we leverage noisy ambiguous sequence calls to understand how the fixation of drug resistance mutations affects diversity. We find that, across all sequences, the presence of drug resistance mutations is associated with lower within-patient genetic diversity, marking the occurrence of selective sweeps. Second, we find that the extent of diversity reduction associated with drug resistance mutations varies with the clinical efficacy of the treatment - efficacious drug treatments with low rates of virologic failure (such as NNRTI-based and boosted PI-based regimens) show strong reductions in diversity associated with each additional resistance mutation, a pattern more consistent with hard selective sweeps, whereas treatments that fail more often (such as regimens based only on NRTIs) show no reduction in diversity, a pattern consistent with soft selective sweeps. Although our results do not explain mechanistically how efficacious treatments lead to harder sweeps of drug resistance mutations, they suggest a more general principle: a lower rate of the production of adaptive mutations should be accompanied by harder sweeps.

## 2 Results

### 2.1 Sequences and patients

We collected sequences of reverse transcriptase and/or protease genes from 6,717 patients from the Stanford HIV Drug Resistance Database [15]. The sequences come from 120 different studies that were performed between 1989 and 2013. The 6,717 patients represent all individuals in the database who were treated with exactly one drug regimen, although this regimen could comprise a combination therapy of multiple drugs (summary in Figure S1). The patients’ viral population was sequenced after at least some period of treatment, although treatment may or may not have ceased at the time of sequencing and treatment may or may not have failed. This virus from a patient was amplified via PCR, sequenced using the Sanger method and then was reported to the database as a single nucleotide sequence. We call this dataset the **D-PCR dataset**.

All 6,717 patients received some type of therapy (between 1 and 4 drugs), with the majority (77%) receiving a regimen of three drugs. Nearly all patients received one or two nucleoside reverse transcriptase inhibitors (NRTI), paired with either a non-nucleoside reverse transcriptase inhibitor (NNRTI) or a protease inhibitor (PI), which was in some patients boosted with a low dose of ritonavir. HIV subtypes were varied, with the majority being B (36%), C (34%) or CRF01_AE (13%). None of the remaining subtypes contributed more than 5% of the total sample.

An additional dataset, which we call the **clonal dataset**, consisted of 11,653 sequences from 740 patients with multiple sequences per patient isolated through clonal amplification and Sanger sequencing. The clonal dataset used only for validation purposes.

### 2.2 Ambiguous calls are a good proxy for genetic diversity

We are interested in the effect of drug resistance evolution on within-patient genetic diversity, but in our main dataset, we only have one sequence per patient. To use this large dataset for our purposes, we therefore use ambiguous nucleotide calls as a proxy for within-patient genetic diversity. Although results from previous studies suggest that this approach is valid [17, 21], we independently validate this measure through comparing the D-PCR and clonal datasets. Using the clonal sequences, within-patient diversity (*π*) can be computed directly, giving an estimate of genetic diversity per site that doesn’t rely on ambiguous calls. We compared the proportion of sequences with ambiguous calls at a site in the D-PCR dataset to the within-patient diversity (*π*) at that site in the clonal dataset (see methods). We find that clonal within-host nucleotide diversity has a high positive correlation with the percentage of nucleotide calls ambiguous in the D-PCR dataset (*r* = 0.91, *p <* 2.2 *×* 10^−16^, Figure S2A). A similar pattern holds at the amino acid level (*r* = 0.85, *p <* 2.2 × 10^−16^, Figure S2B). We therefore conclude that ambiguous calls are a good proxy for within-patient genetic diversity.

### 2.3 Drug resistance mutations (DRMs) lower within-patient diversity

We ask whether across all sequences, the presence of a drug resistance mutation (DRM) is associated with lower within-patient diversity, the classical signature of a selective sweep [9]. For each sequence, we therefore count the number of DRMs present that are relevant for the treatment the patient was taking [22] (i.e., a mutation that confers resistance only to a particular class of drugs was counted as a DRM only if the patient was actually being treated with that drug, see *methods: sequence processing* for more information).

First, for the most common reverse transcriptase and protease DRMs, we compare sequences that have exactly one DRM with sequences that have zero DRMs (i.e., ancestral state at all possible DRM sites). We plot the difference in within-patient diversity between the two groups with 95% confidence intervals in Figure 2A. Among reverse transcriptase and protease DRMs, sequences with the DRM have lower diversity than those with the ancestral state in 14 of 16 cases, with 7 of the 16 being significantly lower at the 95% confidence level. This reduction in diversity is consistent with expectations after a selective sweep [9].

**Figure 2:**
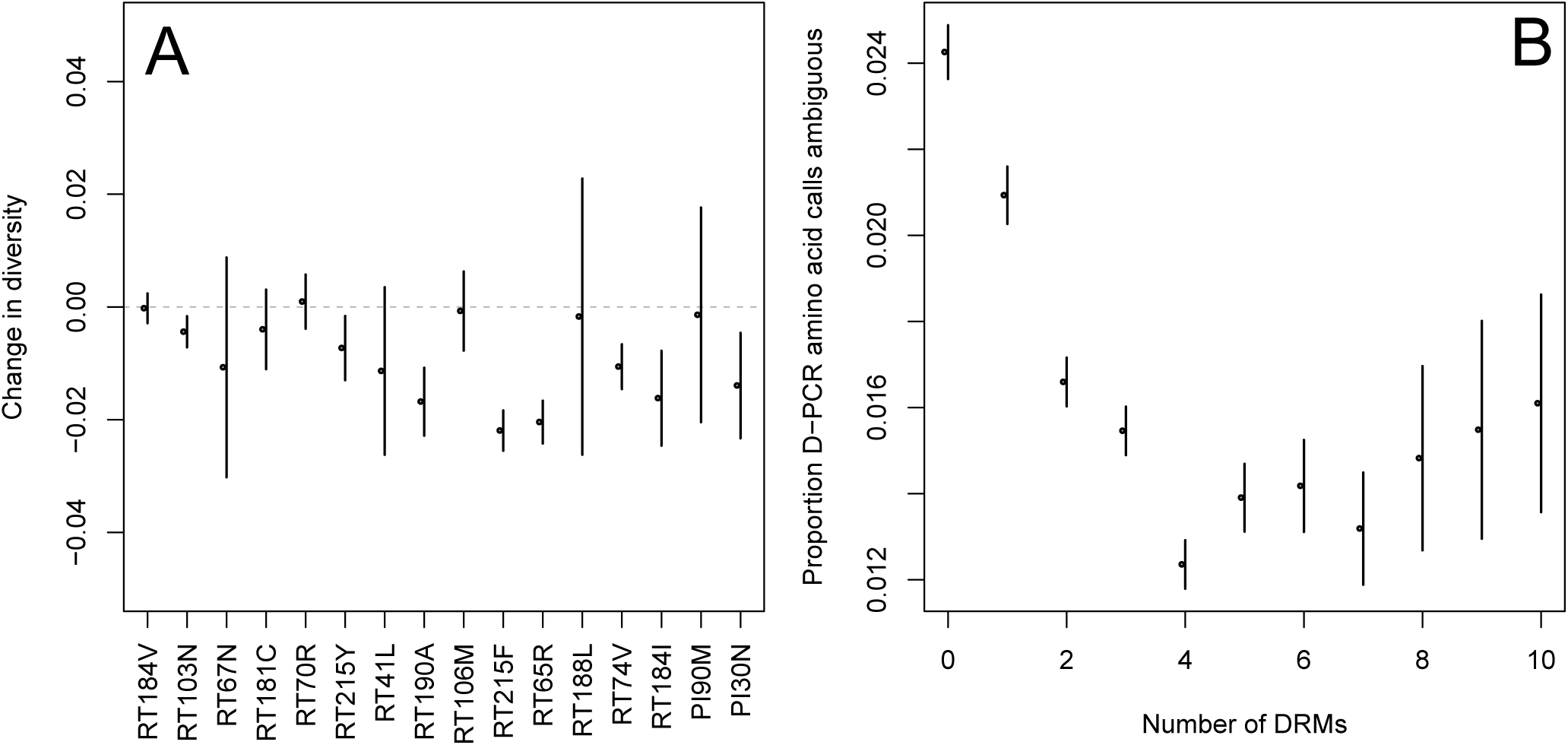
Effect of DRMs on sequence diversity. **(A)** For the most common reverse transcriptase and protease mutations, 95% confidence intervals are drawn for the difference in diversity associated with a single derived mutation. For each DRM, the mean diversity among patients with a fixed ancestral state at the focal locus is compared to those patients with that fixed non-ambiguous DRM. All sequences have no additional DRMs. All DRMs occurring at least 5 times with these specifications are included. **(B)** The effect of multiple DRMs on diversity is shown as the average diversity level of sequences decreases conditional on number of fixed drug resistance mutations present. Means ± SE are plotted among all patients in the D-PCR dataset.

Second, we looked at the effect of multiple DRMs on within-patient diversity, as we hypothesize that multiple fixed DRMs may decrease diversity even more than a single DRM. This could result from sequential selective sweeps of single DRMs each reducing diversity, or from a single selective sweep that fixes multiple DRMs. Indeed, we find that for sequences that have between 0 and 4 DRMs, additional DRMs are associated with reduced genetic diversity (Figure 2B, *p*-value for *t*-test between diversity among sequences with 0 versus *≥*1 is 2.8 *×* 10^−4^, between 1 and *≥*2 is 7.6 *×* 10^−7^, between 2 and *≥* 3 is 0.16, between 3 and *≥* 4 is 1.0 *×* 10^−4^). After 4 DRMs, subsequent DRMs do not significantly change diversity further. The observed pattern of DRMs associated with reduced diversity is mainly driven by the patients receiving NNRTI or boosted PI-based treatments, as can be seen when separating the above analysis by drug treatment category (Figure S3CD). Among patients treated with NRTIs alone or with unboosted PIs, this pattern is much less clear (Figure S3AB). The observed pattern holds across the each of the most common subtypes separately (Figure S4).

### 2.4 Clinical efficacy of anti-retroviral regimens

We have now shown that in general, each additional DRM is associated with reduced diversity, which is consistent with expectations of selective sweeps. We want to test how this effect depends on clinical efficacy of the treatment a patient is on. For the most common drugs in our dataset, we assess clinical drug treatment efficacy categorically and quantitatively.

As a categorical approach, we separated regimens based on general clinical HIV-treatment recommendations where NNRTI-based treatments are preferred to NRTI-based treatments, and treatments based on ritonavir-boosted PIs (PI/r) are preferred to treatments based on unboosted PIs. These more and less efficacious groupings were the basis of comparisons in our parametric approach described below.

To measure efficacy quantitatively, we conducted a literature search to determine the percentage of patients who have remain virologically suppressed after one year of treatment (see *materials and methods, Table S3*) for 21 different treatments with at least 50 sequences per treatment in our D-PCR dataset (see description of **abundant treatment dataset** in *materials and methods: model fitting* for more information.) This quantitative measure was used as the basis for our non-parametric approach described below.

The two measures correspond well and clinical treatment efficacy ranged widely, from very low efficacy (5% of patients virologically suppressed after one year of treatment on AZT monotherapy) to very high efficacy (100% of patients virologically suppressed after one year of treatment on 3TC+AZT+LPV/r) (Figure 3AB).

### 2.5 High treatment efficacy associated with stronger diversity reduction

We hypothesize that efficacious treatments (such as those containing an NNRTI or boosted PI) likely make adaptation in viral populations limited by the generation of mutations and these populations should thus experience harder selective sweeps leading to a sharp reduction in diversity accompanying each additional DRM. Less efficacious treatments on the other hand (such as those containing only NRTIs or unboosted PIs) likely allow replication of fairly large HIV populations so that adaptation is not limited by the generation of mutations. They should thus experience soft selective sweeps and little or no reduction of diversity with each additional DRM. Below we test this hypothesis by assessing the reduction of diversity associated with the presence of a DRM among treatments that vary in clinical efficacy.

To determine whether the effect of DRMs on within-patient diversity depends on clinical treatment efficacy, we first fit generalized linear models (GLMs) separately for each of 29 treatments (see Figure S1) using the year and the number of DRMs to predict diversity. For each treatment we report the effect of the number of DRMs on diversity as fit by the GLM as ∆_*DRM*_ (3CD). We included only treatments that had 50 sequences, a sufficient number of observed patients with different numbers of DRMs (see description of **abundant treatment dataset** in *materials and methods: model fitting* for more information) and use only sequences with at most 4 DRMs. The same analysis with all sequences is repeated in the supplement (Fig S5) and yields qualitatively similar results. Lower ∆_*DRM*_ values correspond to a bigger decrease in diversity associated with a ∆_*DRM*_ - a pattern more consistent with hard selective sweeps.

We find that most of our ∆_*DRM*_ estimates are qualitatively consistent with expectations: efficacious treatments have lower ∆_*DRM*_ values than less efficacious treatments. Most NNRTI-based treatments are associated with a reduction of diversity per DRM (∆_*DRM*_). In 10 of 11 NNRTI-based treatment regimens, ∆_*DRM*_ is significantly below 0 (Figure 3C). This pattern suggests the presence of hard sweeps. Less efficacious treatments containing only NRTIs were generally associated with a smaller or no reduction in diversity per DRM, although 4 of the 7 treatments with 2 or 3 NRTIs also had a ∆_*DRM*_ value significantly below 0. In some cases, such as DDI or AZT monotherapy, there was even an increase of diversity associated with DRMs (a significantly positive ∆_*DRM*_ value, see figure 3C). This pattern is suggestive of soft sweeps.

**Figure 3:**
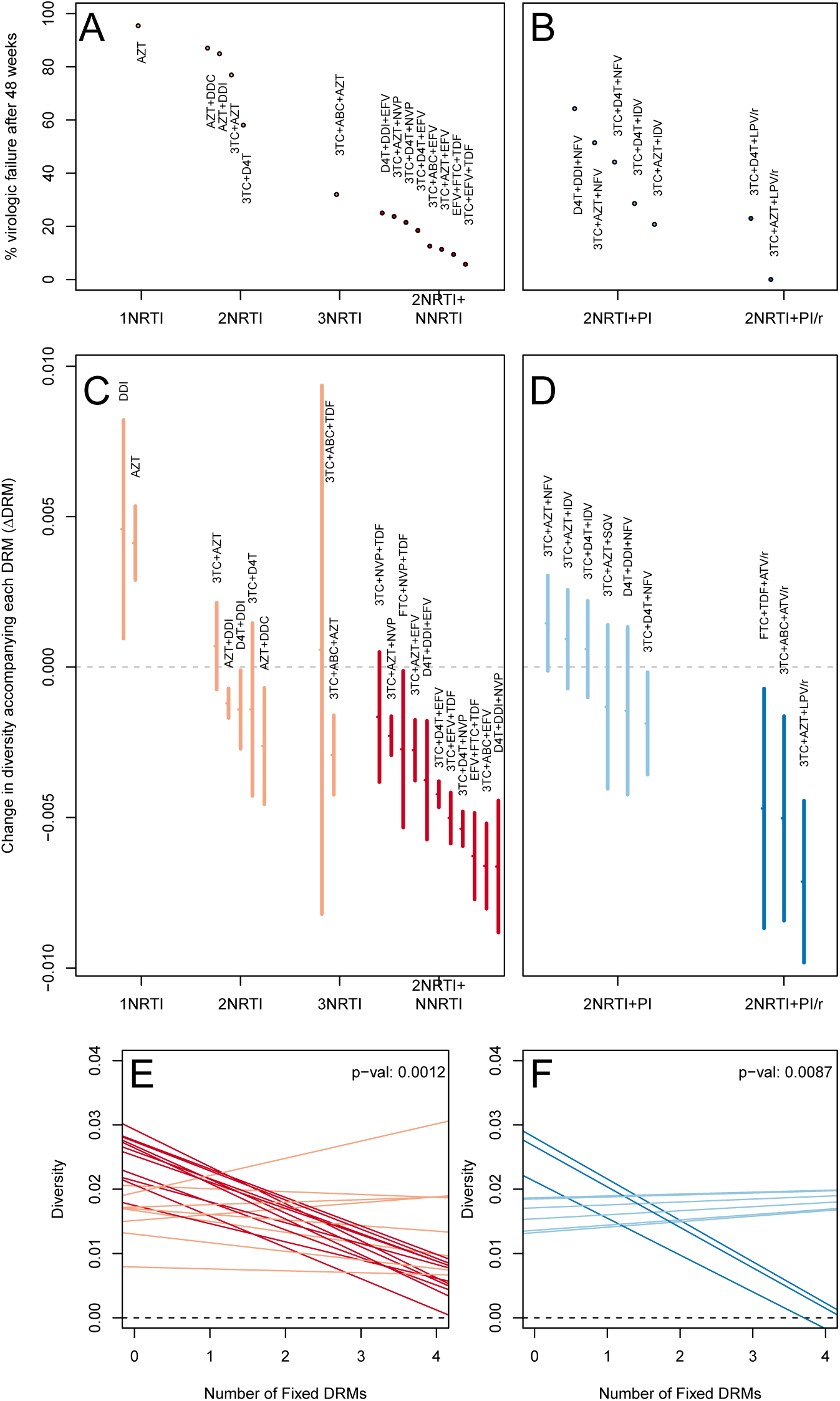
Drug resistance mutations are correlated with diversity reduction differently in different types of treatments. Treatment efficacy from literature review (% of patients with virologic suppression after ̰ 48 weeks) showed positive correspondence with clinical recommendation among RTI regimens (**A**) and PI+RTI regimens (**B**). ∆_*DRM*_ ±SE lower among the more efficacious and clinically recommended treatments among RTI treatments (**C**) and RTI+PI treatments (**D**). Mixed effect model shows significantly different slopes for NNRTI treatments versus NRTI treatments (**E**) and PI/r treatments versus PI treatments (**F**). Each line in (**EF**) represents the fitted decay in diversity with each DRM for a different treatment from the full mixed effects model and p-value labeling indicates the difference between the plotted full model and the null not fitting slopes separately for treatment groups.

We found a slightly positive value of ∆_*DRM*_ for the NRTI-based regimen 3TC+ABC+TDF, a treatment which is known to often lead to rapid treatment failure [23, 24]. Additionally, we found that the regimen 3TC+NVP+TDF had the least negative ∆_*DRM*_ value among the NNRTI-based treatments, which is consistent with findings from Tang et al suggesting treatment inferiority [25]. Among the most negative ∆_*DRM*_ values of the NNRTI-based treatments were the treatments 3TC+ABC+EFV and EFV+FTC+TDF, which have long been on the list of recommended treatments in the USA [26] (until their recent replacement with INSTI-based treatments which are not in our dataset). Somewhat surprisingly, the most negative ∆_*DRM*_ value was associated with D4T+DDI+NVP, which is not a recommended treatment.

Among treatments containing PIs, all three of the efficacious boosted-PI treatments had ∆_*DRM*_ significantly below 0 (Figure 3D). The less efficacious unboosted PI treatments had ∆_*DRM*_ values closer to 0, although 1 of 5 unboosted treatments (3TC+D4T+NFV) also had a ∆_*DRM*_ value significantly below 0. As expected, we find that the LPV/r treatment in our dataset has a much lower ∆_*DRM*_ value than the NFV treatments, consistent with the recommendations that LPV/r is preferable to NFV in relation to drug resistance prevention [27].

To quantify and further test the observation that more clinically efficacious treatments lead to the greater diversity reduction per fixed DRM, we use two primary approaches, one parametric and the other non-parametric.

#### Parametric approach

This analysis is done separately for non-PI-based treatments and PI-based treatments. For each of these two broad classes, we separate treatments in to high and low efficacy groups, so that we first compare NRTI-based treatments with NNRTI-based treatments and then we compare non-boosted with boosted PI-based treatments. For each of the two broad classes of treatment, we fit a mixed effects model to separately fit ∆_*DRM*_ slopes for high and low efficacy treatments, allowing us to quantify the difference between the two treatment types. Using an ANOVA to compare this model to one which does not distinguish between treatment types, we measure the extent to which fitting the types separately improves model fit (see *materials and methods: model fitting*).

We find that among sequences from patients receiving highly efficacious treatments with NNRTIs each fixed DRM is associated with an additional 19% reduction in diversity as compared to poplations with no DRMs, whereas among sequences from patients receiving less efficacious treatments with only NRTIs, each fixed DRM is associated with an additional 0.6% reduction in diversity compared to populations with no DRMs. (Table S1). The relative effect of these ∆_*DRM*_ coefficients can be seen in Figure 3E, where dark red lines are model fits for efficacious NNRTI treatments and light red lines are model fits for less efficacious NRTI treatments. Further, in comparing the relative fits of this model to a null model not separating the more and less efficacious treatments, we find that the model separating out the two treatment types significantly improves the fit to the data (ANOVA, *p* = 1.2 *×* 10^−3^) suggesting that the fixation of a DRM is associated with reduced diversity much more when treatments are efficacious.

In an analogous analysis limiting our dataset to PI-based treatments, we found that sequences from efficacious treatments based on boosted PIs showed an additional reduction in diversity of 24% with each fixed DRM as compared to populations with no DRMs, while sequences from less efficacious treatments based on unboosted PIs showed a slight increase in diversity by 3% with each fixed DRM (Table S1). The relative effect of these ∆_*DRM*_ coefficients can be seen in Figure 3F, where dark blue lines are model fits for efficacious PI/r treatments and light blue lines are model fits for less efficacious unboosted PI treatments. The ANOVA comparing the models suggests that fitting the effect of DRMs on diversity separately for treatments with boosted and unboosted PIs results in a better fit of the data (ANOVA, *p* = 8.7 *×* 10^−3^). In both treatment classes, we find that the fixation of a DRM under a more efficacious treatment leads to a much stronger reduction of diversity than the fixation of a DRM under less efficacious treatments.

#### Non-parametric approach

As a second, non-parametric approach, we did a test with the ability to compare the effects of all treatments (RTIs and PIs) directly, we fit a linear regression among the 21 treatments for which we had efficacy information predicting the reduction of diversity per DRM from treatment efficacy. We find that in a stretch of 1000 basepairs, a 10% increase in treatment efficacy is associated with 2 fewer ambiguous nucleotide calls among patients with 3 DRMs as compared to those with 0 DRMs. This means that patients given treatments with 30% efficacy have approximately the same amount of diversity whether or not they have 0 or 3 DRMs, but patients on treatments with 80% efficacy have 10 fewer ambiguous calls over a 1000 basepair region of reverse transcriptase if they have 3 DRMs as compared to 0. In terms of ∆_*DRM*_, a 10% increase in treatment efficacy is associated with a −6.68 *×* 10^−04^ decrease.

Next, we do a randomization test to determine whether the linear regression could be the result of year as a confounding factor (see: Model Fitting). We find that the fitted slope in the real data was lower than 98.17% of our randomized slopes, suggesting that our observed association between efficacy and diversity reduction cannot be explained by the of sampling year alone. However, we did find that the mean of our randomized regression slopes was centered below 0, indicating that the confounding factor of year was responsible for some, but not all of our signal. In examining our mixed effects models, year had a smaller effect on diversity than the number of drug resistance mutations (Table S1).

## 3 Discussion

Treatment for HIV-1 represents an enormous success of modern medicine. Whereas early antiretroviral treatments were associated with fast evolution of drug resistance and high rates of treatment failure, there are currently many combinations of drugs that are successful at keeping HIV-1 at low or undetectable levels for many years, preventing the evolution of resistance and the progression to AIDS. Indeed, the evolution of drug resistance has become fairly uncommon [28]. In this paper, we’ve shown that this shift from fast evolution of drug resistance to slow has been accompanied by a corresponding shift from soft selective sweeps (in which the same DRM occurs on multiple genetic backgrounds) to hard selective sweeps (in which a DRM occurs only a single time). This suggests that modern treatments have brought HIV into a regime where the viral population must wait until the correct mutation or combination of mutations is generated. This also means that for any given patient the acquisition of drug resistance has become at least partly an unlucky occurrence, in sharp contrast to the early days of HIV treatment in which all patients predictably failed treatment. Harder sweeps within well-treated patients are also consistent with the overall decrease in the rate of resistance.

We want to study how the evolutionary dynamics of selective sweeps have changed in the evolution of drug resistance over the past two and a half decades. Because it would be unethical to give subpar treatment to HIV infected patients, we can only investigate this question using historical data. The only type of data that is available for a wide range of treatments and time points is Sanger sequencing data, due to its importance in HIV research and diagnostic testing. Although we often have only one sequence per patient, it is important to note that there is still information present in these sequences about diversity and selective sweeps. First, we used ambiguous nucleotide calls as a proxy for genetic diversity. Although we are not the first study to do so ([17, 18, 18–20]), as far as we are aware, we are the first to use clonal sequences to validate its accuracy as a measure. Second, we can use the number of fixed drug resistance mutations to determine how much adaptation to treatment has taken place. We used a fairly conservative list of DRMs very unlikely to fix in the absence of the drug [29–31]. Therefore, DRMs must have fixed as a result of strong positive selective pressure imposed by the drug, and are indicative of recent selective sweeps. We can then look at the correlation between the number of drug resistance mutations and genetic diversity and see if that relationship has changed across treatments and time. Because this approach relies only on widely-available Sanger sequences, we were able to compare 29 different treatments, from AZT monotherapy to treatments based on boosted PIs, sampled across more than two decades (1989 - 2013).

Examining the relationship between DRMs and diversity recapitulates expected results: we first find that across the entire dataset, sequences with a single DRM have lower genetic diversity than sequences without any DRMs. This result confirms a finding from a previous much smaller study looking at patients on NNRTI based treatments ([32]). In addition, we find that having more DRMs is associated with a greater reduction in diversity. This pattern could be generated by successive selective sweeps, in which DRMs are fixed one by one and each selective sweep lowers the diversity further. Alternatively, multiple DRMs may have fixed simultaneously in a single selective sweep.

The key result of this paper (illustrated in Figure 1) is that drug resistance mutations are associated with reduced diversity in patients on efficacious treatments, whereas this pattern is not seen among patients on older treatments with low clinical efficacy. For example, among patients given treatments with 30% efficacy, sequences with 3 DRMs are predicted to have marginically fewer ambiguous calls as those with 0 DRMs (0.5 fewer ambiguous reads over 1000 bases). In contrast, among those patients given treatments with 80% efficacy, sequences with 3 DRMs are predicted to have 10 fewer ambiguous calls than those with 0 DRMs over 1000 bases, a substantial decrease in genetic diversity. Thus, the higher the treatment efficacy, the more DRMs are associated with low genetic diversity. This is consistent with drug resistance evolution dominated by soft selective sweeps when failure rates were high, transitioning over time to evolution dominated by hard selective sweeps as treatments improved and failure rates became much lower. Clinically efficacious treatments are thus characterized by a more frequent occurrence of hard selective sweeps.

It is of interest to compare our new results to a previous study by one of us, Pennings *et al* [32]. Sequences in that study came from patients who were mostly treated with EFV + IDV, a combination that never became common and is not represented here, but which has an efficacy of 75% [33]. The study examined selective sweeps in those patients by looking at the fixation of a particular DRM, at amino acid 103 in RT, which changes from Lysine (K, wild type) to Asparagine (N, resistant). K103N is special because it can be caused by two different mutations, as the wild type codon AAA can mutate to AAT or AAC, both of which encode Asparagine. When focusing on patients whose virus acquired the K103N mutation, the study found that in some patients, both the AAT and AAC codons were found (which is clear evidence that a soft sweep has happened, see Figure 1 in the original paper), whereas in other cases only one of the two was there (which suggests that a hard sweep may have happened, see Figure 2 in the original paper). Because of the detailed data available for these patients, it was shown that both soft and hard sweeps were occurring almost equally often. Placing the 75% efficacy of EFV+IDV in the context of our above results, this is also what we would have predicted. Now we know that this result (hard and soft selective sweeps occur) is not something that will be generally true for HIV, but rather it is a function of the efficacy of the treatment. Had Pennings *et al* had data from a much worse or much better treatment, they may have concluded that hard sweeps or soft sweeps were the rule in HIV.

The transition to highly efficacious treatments and hard selective sweeps was not abrupt. As visible in Figure 1, treatment efficacies and ∆_*DRM*_ do not cluster into distinct groups based solely on the number and type of component drugs. The incremental changes in efficacy and the evolutionary dynamics are worth noting because a simplified narrative sometimes suggests that solving the drug resistance problem in HIV was achieved simply by using three drugs instead of two [34]. According to this narrative, HIV can always easily evolve resistance when treatment is with one or two drugs, but it is virtually impossible for the virus to become resistant to three drugs. In truth, particular combinations seem to lead to more favorable evolutionary dynamics for patients (as seen in Figure 3AB).

Many potential mechanisms could drive the observation that more efficacious drugs drive hard sweeps in wtihin-patient populations of HIV-1. Better drugs may allow for a faster collapse of population size, decreasing the probability that one or more ‘escape’ DRMs occurs ([35], though see [36]). Alternatively, suppressed HIV populations may continue replicating at small numbers, and better drugs may cause this replicating population to be smaller than among patients given inferior drugs. Similarly, a treated patient may retain a reservoir of HIV unreachable by the treatment, and better drugs may make this reservoir smaller. If newer drugs have fewer side effects and therefore improved adherence among patients, this too could result in a smaller within-patient population size among patients treated with better drugs, and contribute to the decreased production of resistant genotypes. The effect could be driven by standing genetic variation, if worse drugs would allow preexisting mutations to establish, whereas better drugs make this less likely, for example, if a pre-existing mutation could only establish if it occurred in a small and specific compartment [36]. Finally, more efficacious drugs may simply require more mutations on the same background (i.e., a higher genetic barrier to resistance [25]), so the effective mutation rate to a fully resistant genotype is lower. Our data does not give us sufficient resolution to distinguish between these hypotheses, and the true dynamics may be a combination of all of these factors. However, all these potential mechanisms work to reduce the rate of production of resistant genotypes and the principle is more general: if the probability of acquiring resistance is low, rare resistance should be generated by hard selective sweeps.

There are a number of caveats to our analysis. First of all, this is a cross sectional study, so we cannot establish causation. Second, because our data were collected from a heterogeneous variety of sources over a substantial time period, it is possible that sampling issues may be influencing our results. However, given that our primary measure to understand the effect of the fixation of a drug resistance mutation is an internal comparison with other patients on the same treatment, we do not believe that our results could be generated solely by data heterogeneity. Further, that our results are robust to different measurements and filtering approaches suggests DRMs do indeed sweep differently in HIV populations subject to more or less efficacious treatments. Thirdly, we are limited by the discovered DRMs. The HIV genome is extremely well annotated for sites related to drug resistance, but of course, it this annotation not perfect. If we see a no DRMs fixed, a mutation unknown to us may have fixed. This may be especially true for PI resistance, which is not as well understood as resistance to RTIs. ∆_*DRM*_ could thus appear less negative if a mutation elsewhere in the genome has swept and lowered diversity, but we count zero DRMs within protease. This may make our documented signal conservative. Alternatively, if one DRM is observed in protease, but a second DRM occurred elsewhere and is therefore not observed in our sample, ∆_*DRM*_ may appear more negative, because the one DRM category actually has the diversity reduction of two DRMs. While drug resistance to PIs is complex, this behavior should not apply differently between regimens in a systematic way, and is therefore unlikely to drive the observed results.

Drug resistance evolution is no longer the threat to HIV patients it once was and treatments exist that almost never lead to resistance. However, our observation that efficacious drug combinations lead to hard selective sweeps could be useful for improving treatments for other pathogens. Even among relatively small samples, if patients who acquire drug resistance mutations do not have a significant decrease in diversity relative to those who did not acquire drug resistance mutations, this might suggest that the fixing mutations are occurring via soft selective sweeps, and the treatment brings patients into a dangerous regime for drug resistance. Alternatively, if patients acquire drug resistance mutations, but those patients also have very little diversity, it may be that this is a safer treatment and that the emergence and fixation of drug resistance mutation was a relatively uncommon occurrence.

Looking at changes in diversity following a sweep in order to assess the mode of adaptation could be particularly well-suited to looking at evolution of cancer. While single cell sequences isolated from tumors have yielded promising insights about evolutionary dynamics, the process is invasive and relatively difficult. Sequencing tumor-free cancerous cells circulating in the blood provides less information, but can be done serially and provides a good measure of tumor heterogeneity. Applying a method such as ours to monitor changes in cell-free DNA diversity over time may allow us to determine if certain treatments reproducibly lead to soft sweeps.

Comparing treatment efficacy with the occurrence of soft and hard selective sweeps may also provide supplementary information about additional risk factors among patients. In the case of HIV, we find that high efficacy is associated with hard selective sweeps, which suggests that the the virus has a hard time evolving drug resistance, and the patients in whom resistance evolved are merely the unlucky ones. However, there may be cases where failure is rare, but associated with soft selective sweeps. Such a situation thus reflects a discordance between what happens within patients and what we see at a population level. This, in turn, may be indicative of a behavioral, genetic or virologic difference among the groups of patients and efforts should be made to find out how failing patients are different from non-failing patients.

In conclusion, we find that the study of diversity in viral populations with resistance can show differences in evolutionary pathways of adapting pathogenic populations and provides a concrete example of how population genetics theory can make substantive predictions about medically relevant problems. Next generation and single molecule sequencing have the capacity to bring much more precision in determining the dynamics of within-patient populations. However, we also urge researchers and clinicians to report more information concerning the diversity of pathogen populations, even in the form of minor allele frequency cut offs for calling ambiguous calls or raw sequencing data, as this might allow new insight from data that might be otherwise overlooked.

## 4 Materials and Methods

### 4.1 Data Collection & Filtering

#### Direct PCR (D-PCR) dataset

We collected one consensus sequence of the HIV-1 reverse transcriptase gene per patient from 6,717 patients across 120 different studies from the Stanford HIV Drug Resistance Database [15]. These patients represent all individuals in the database with HIV populations which were treated with exactly one drug regimen and that had an associated reverse transcriptase sequence. Protease sequences were also recorded, when available (5163 sequences).

Nearly all patients received treatment that included one or more nucleoside reverse transcriptase inhibitors (NRTI). In many patients, the NRTIs were paired with either a non-nucleoside reverse transcriptase inhibitor (NNRTI) or a protease inhibitor (PI), which was in some patients boosted with a low dose of ritonavir, a second PI, in order to boost drug levels and improve efficacy. Among patients receiving protease inhibitors (PIs), only patients with both protease and reverse transcriptase sequences were included. All sequence information in the HIV database is recorded as an aligned consensus nucleotide sequence. The data are taken across many studies, each with their own procedures and cutoffs for calling positions ambiguous. No electropherogram data was available and the ambiguous calls reported were taken as submitted to the database. The sequences from these patients were labeled the **direct PCR (D-PCR)** dataset.

#### Abundant treatment dataset

For each sequence we counted the number of relevant drug resistance mutations (DRMs, see below). For a portion of the analysis, we rely on examining many HIV-1 sequences from patients being treated with the same drug regimen. Because we measure the association between the number of DRMs and genetic diversity by examining a slope (∆_*DRM*_), our signal could be heavily influenced by single patients if the distribution of patients with a certain number of DRMs within a given treatment is narrow (e.g., a treatment has almost only patients with 0 DRMs). To ensure that that our observed signal is not driven by these cases, we exclude treatments if among the sequences from a treatment we do not have at least three sequences within three of the DRM categories (0, 1, 2, 3 or 4 DRMs). Distributions of number of DRMs per sequence are shown in the supplement for all treatments (Figure S5). This restriction yielded a dataset with 5147 sequences from 29 treatments and was termed the **abundant treatment** dataset.

The breakdown of the D-PCR dataset and the abundant treatment dataset by the kind of treatment a patient got (protease inhibitor based treatment and reverse transcriptase inhibitor based treatment) is listed in Figure S1.

#### Clonal dataset

We supplemented our analysis with an additional dataset of patients from which multiple HIV-1 strains were sampled. This dataset comprised 10,235 sequences, but from a relatively small number of patients (*n* = 174) with much less variety in their given treatments (the mean number of sequences per patient was 58 with an interquartile range of 16 to 64). These sequences were derived from taking a within-patient population of HIV-1 and attempting to isolate, clone and sequence single strains. We used these reads (termed the “clonal dataset”) to validate ambiguous calls as a within-patient diversity measure.

### 4.2 Sequence Processing

Sequences from reverse transcriptase and protease were analyzed to determine the number of ambiguous calls and the number of drug resistance mutations per sequence.

We called a nucleotide non-ambiguous if it read A, T, C or G, and grouped lowercase (and less confident) a, t, c and g calls with their capital counterpart. Nucleotides called as W, S, M, K, R, Y (ambiguity between two nucleotides) and B, D, H, V (ambiguity between three nucleotides) and their lowercase counterparts were included as ambiguous calls. Ns and Xs (indicating no information about the identity of the position) were excluded.

We also examined ambiguities on the amino acid level by using nucleotide level information. If, for example, a nucleotide triplet was recorded as AAM, where M indicates an adenine/cytosine ambiguity, the amino acid at that position was ambiguous between AAA (Lysine, K) and AAC (Asparagine, N). The amino acid for that position would then be recorded K/N. All ambiguous calls at the nucleotide level were translated into ambiguous calls at the amino acid level, including if the ambiguous call reflected synonymous encodings (i.e., AAA and AAG are both Lysine, and the amino acid would be encoded K/K). The number of ambiguous amino acids was recorded for each sequence and was normalized by the length of the amino acid sequence.

We determine whether mutations are associated with drug resistance based on the 2009 update of *DRMs for the surveillance of transmitted HIV-1 drug resistance* adopted by the World Health Organization [22], which lists drug resistance mutations that are indicative of selective pressure. For each patient, we determine the number of mutations that confer drug resistance at the amino acid level to any of the classes of drugs the patient is receiving (i.e., NRTI, NNRTI, PI). These drug resistance mutations are counted as fixed if they are either non-ambiguously the resistant type, or they are an ambiguous call with all possible states as the resistant type (i.e., N/N in the example above).

5163 patients in the D-PCR dataset had a sequenced protease gene available in the database (77% of patients). Not all patients had entire reverse transcriptase and protease genes sequenced, but only 1% of sequences had fewer than 500 basepairs sequenced of 1680 possible in reverse transcriptase and only 1% of available protease sequences had fewer than 210 of 300 bases in protease. We normalized the number of ambiguous calls by sequence length to calculate the probability of observing an ambiguous call per amino acid.

### 4.3 Validation of ambiguous calls as genetic diversity measure

In order to validate the appropriateness of ambiguous calls as a proxy for genetic diversity, we computed diversity using an ambiguous call measure and compared it to a diversity measure that did not use the ambiguous call data. For each site, we computed genetic diversity using the ambiguous calls as the proportion of all D-PCR HIV-1 sequences that had an ambiguous basepair call at that site. This approximates the percentage of patients with within-patient diversity by site.

In order to test if this measure of diversity based on ambiguous calls correlates well with other measures of diversity that don’t depend on ambiguous calls, we used the clonal dataset to compute the average sitewise *π*. For a site with nucleotides *A*, *T*, *C* and *G* at within-patient frequencies *p*_*A*_, *p*_*T*_, *p*_*C*_ and *p*_*G*_, *π* is computed as

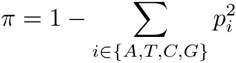

*π* is equal to zero when every sequence has the same basepair call (i.e., all As) and is maximized when multiple categories are at intermediate frequencies (an even split between A,T,C and G). Within-patient diversity was measured by first computing *π* at each site separately within each patient and then averaging over all patients. The *π* calculation at a site for a particular patient was only included if the patient had at least two sequences calling non-N identity at that site. We computed diversity in the same way at the amino acid level to validate that our signals persisted when looking at codons.

Because the drug regimens systematically changed over time (Figure S6), it is essential to ensure that our estimates of diversity are not confounded by the changes in sequencing practices over the same time. While all samples were Sanger sequenced, we do find that the number of recorded ambiguous calls increased over time (Figure S6C), possibly because the cut-off of calling a read as ambiguous became lower, however the effect was not significant (*p* = 0.16, linear regression with year predicting the proportion of ambiguous calls).

### 4.4 Clinical efficacy of antiretroviral treatments

We expect that clinical efficacy of an HIV treatment affects the probability of the virus undergoing a soft or hard selective sweep. All HIV treatment regimens that occurred at least 50 time in the D-PCR dataset were evaluated for treatment efficacy based on a literature review. As a measure for treatment efficacy, we recorded the proportion of patients whose treatment was still successful after a year of treatment, as indicated by a viral load of *≤* 50 copies of HIV-1 RNA/mL or less after 48 or 52 weeks in an on-treatment analysis. Our literature review was mostly based on the papers reviewed in Lee et al [28]. Because this review did not include review information for several older treatment regimens, we supplemented our analysis with additional studies. A full description of how clinical treatment efficacy was calculated by study can be found in the supplement (Supplement: Determining treatment efficacy, Table S2). A second researcher randomly chose 5 studies and independently followed the protocol to determine treatment efficacy for these studies, providing confirmation of our method. Because we believe the thus collected information may be useful to other researchers, we provide our estimates in the supplement (Table S3).

### 4.5 Quantifying the relationship between clinical efficacy and diversity reduction

For each treatment, we estimate the relationship between the number of DRMs and genetic diversity (delta DRM) by fitting a generalized linear model. As time may be a confounding factor in our analysis, year of sequence collection (reindexed so 1989 maps on to 0) was included as a covariate:

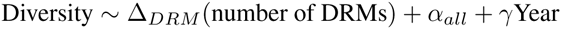

Sequences with more than four DRMs were excluded. We performed this truncation both because the data becomes non-monotonic after four DRMs (see Figure 2), but also because different treatments had more or fewer patients with large numbers of DRMs which would render results from a linear model incomparable. Not excluding these sequences produced qualitatively similar results (Figure S7). To quantify the effect of treatment efficacy on delta DRM, we used both a parametric and a non-parametric approach.

#### Parametric approach

To compare how the effect of DRMs on genetic diversity varied between two groups of treatments, we fit mixed effects models in R [37] using the package mle4 [38] including all sequences belonging to the 29 treatments that passed our threshold criteria (see above). These models were parametrized to fit separate slopes (delta all and delta e) and intercepts (alpha all and alpha e) for comparisons among the two different types of treatments (NRTIs+NNRTI were compared to NRTIs only; NRTIs+PI were compared with NRTIs+PI/r) in addition to random effects slopes (delta t) and intercepts (alpha t) being fit separately for each treatment regimen within each treatment group. To do this, we used an indicator variable for membership in the higher efficacy group (NNRTI or PI/r groups) and fit the following model:

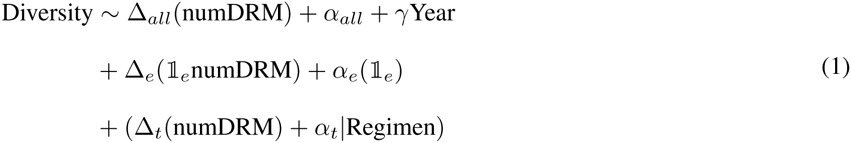

where

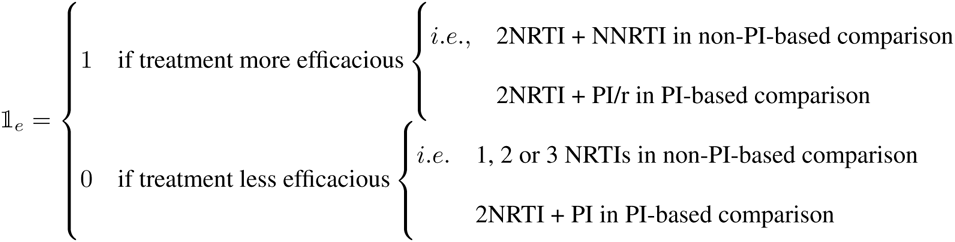

and subscripts *t, e* and *all* correspond to treatment-specific, efficacious group and total sample effects, respectively. *α* terms correspond to intercepts in the model.

The diversity change associated with a DRM for a given treatment is ∆_*DRM*_ = (∆_*all*_ + 𝕀_*e*_∆_*e*_ + ∆_*t*_). Therefore, ignoring treatment specific effects, the overall ∆_*DRM*_ associated with the less efficacious group is ∆_*all*_ and the ∆_*DRM*_ associated with the more efficacious group is ∆_*all*_ + ∆_*e*_. In this way, we can interpret (∆_*e*_) as the diversity change attributable to being in the more efficacious group. The intercepts were similarly fit separately among the two groups to account for differing baseline levels of diversity.

#### Non-parametric approach

Apart from discriminating between two treatment categories, we also tested the association between a continuous measure of treatment efficacy and the change in diversity associated with a DRM on a given treatment (∆_*DRM*_) as a non-parametric approach. We then fit a linear regression between treatment efficacy and the corresponding ∆_*DRM*_ values to quantify the effect of treatment efficacy on the diversity reduction associated with drug resistance mutations. We called the slope of this line *α_dat_*.

To test the significance of *α_dat_*, we resampled our data 10,000 times, randomly assigning HIV population sequences to treatments, but controlling the sampling so that each treatment had the same sample size composed of sequences with the same distribution of years. For each resampled treatment, we found the corresponding simulated ∆_*DRM*,*sim*_ term and refit the linear regression to determine our randomized slope *α_sim_*. These randomized coefficients were then used to assess the significance of the the relationship between diversity and treatment efficacy in our observed data.

## 5 Acknowledgments

The authors thank Alison Hill, Daniel Rosenbloom, Heather Machado, Nandita Garud, Ben Wilson, Emily Ebel, Zoe Assaf, Rajiv McCoy, Richard Neher, Joachim Hermisson and members of the Petrov lab for comments and discussion. SYR and RWS were supported in part by NIH grant R01 AI068581. This material is based upon work supported by the National Science Foundation Graduate Research Fellowship to AFF under Grant No. DGE-114747.

